# Developmental changes in individual alpha frequency: Recording EEG data during public engagement events

**DOI:** 10.1101/2023.01.20.524682

**Authors:** Christopher Turner, Satu Baylan, Martina Bracco, Gabriela Cruz, Simon Hanzal, Marine Keime, Isaac Kuye, Deborah McNeill, Zika Ng, Mircea van der Plas, Manuela Ruzzoli, Gregor Thut, Jelena Trajkovic, Domenica Veniero, Sarah P Wale, Sarah Whear, Gemma Learmonth

## Abstract

Statistical power in cognitive neuroimaging experiments is often very low. Low sample size can reduce the likelihood of detecting real effects (false negatives) and increase the risk of detecting non-existing effects by chance (false positives). Here we document our experience of leveraging a relatively unexplored method of collecting a large sample size for simple electroencephalography (EEG) studies: by recording EEG in the community during public engagement and outreach events. We collected data from 346 participants (189 females, age range 6-76 years) over 6 days, totalling 29 hours, at local science festivals. Alpha activity (6-15 Hz) was filtered from 30 seconds of signal, recorded from a single electrode placed between the occipital midline (Oz) and inion (Iz) while participants rested with their eyes closed. A total of 289 good quality datasets were obtained. Using this community-based approach, we were able to replicate controlled, lab-based findings: IAF increased during childhood, reaching a peak frequency of 10.28 Hz at 28.1 years old, and slowed again in middle and older age. Total alpha power decreased linearly, but the aperiodic-adjusted alpha power did not change over the lifespan. Aperiodic slopes and intercepts were highest in the youngest participants. There were no associations between these EEG indexes and self-reported fatigue, measured by the Multidimensional Fatigue Inventory. Finally, we present a set of important considerations for researchers who wish to collect EEG data within public engagement and outreach environments.

## 1. Introduction

Cognitive neuroscience is primarily a laboratory-based endeavour. Although lab-based neuroimaging experiments are often limited in terms of the ecological validity of the behaviours that are studied (Ladouce et al., 2017), studying participants within the lab offers numerous benefits to the researcher in terms of experimental control. For example, environmental and physiological artifact can be minimised when recording brain activity using electroencephalography (EEG), thereby enhancing the signal-to-noise ratio of the data that is collected. Lab-based studies can also facilitate the application of higher-density electrode arrays, and the completion of long, time-consuming experiments involving many hundreds, if not thousands, of trials. At a logistical level, much of the hardware used in cognitive neuroscience is also expensive, fragile, and not portable, and thus researchers may have little choice but to require participants to visit the lab to answer specific scientific questions.

However, lab-testing is slower particularly in the case of testing specific populations e.g., children, older people, or those with certain clinical diagnoses. As a result, the average sample size in EEG experiments is generally small: Clayson et al., (2019) identified an average sample size of only 21 participants across a random selection of ERP papers in 5 high-impact cognitive neuroscience journals. This under-recruitment is counterproductive, since small effect sizes are common in cognitive neuroscience and large numbers of participants are needed to detect them (Ioannidis, 2005). As a result, Button et al. (2013) estimate that the average statistical power of studies in neuroscience is very low, leading to poor reliability and reproducibility of the reported findings. Many researchers rely on recruiting from the locally available pool of undergraduate students, who have low diversity of age, educational attainment, socio-economic status, race, and ethnicity (Dotson & Duarte, 2020; Henrich et al., 2010). Furthermore, people with disabilities, neurodiversity, mental health issues and even left-hand dominance are often excluded as “atypical”, thereby exacerbating the poor representativeness of the research sample relative to the wider population (Falk et al., 2013). Although many researchers have recently moved towards collecting data in an online context - either as a conscious choice to improve sample size and diversity, or as a necessary response to Covid-19 restrictions - this approach is clearly not a feasible alternative for neuroimaging studies.

Several initiatives have been implemented over recent years to increase sample size and to improve the rigour, reproducibility, and representativeness of EEG research. Open databases, such as the NEMAR gateway (Delorme et al., 2022) and the Healthy Brain Network (Alexander et al., 2017) provide access to large, ready-made collections of EEG data that can be re-analysed, thereby reducing or eliminating the need to record additional data locally. Many journals also now mandate that datasets are made openly available after a manuscript has been accepted for publication (see White et al., 2020, for a discussion of the benefits and challenges of data sharing in neuroimaging research). Secondly, large-scale collaboration networks such as the #EEGManyLabs initiative (Pavlov et al., 2021) and ENIGMA-EEG (Smit et al., 2021) provide frameworks for multiple, geographically distributed labs to pool participants to answer scientific questions, including multi-lab replications of seminal studies. Finally, recent technological advancements in mobile EEG systems have also made it easier to record high quality electrophysiological data in more ecologically valid environments (Gramann et al., 2011). These mobile systems should be seen as an important step forward in bringing cognitive neuroscience out of the lab and into the community, with the potential to also foster improved participant diversity in the data that is collected.

One lesser-explored method of collecting large numbers of EEG datasets within the community is via public engagement and outreach events. In the appropriate environment, a public engagement stall can engage a considerable number and breadth of people from diverse and often poorly-engaged groups. The National Co-ordinating Centre for Public Engagement defines public engagement as “*the myriad of ways in which the activity and benefits of higher education and research can be shared with the public*”. In doing so, they rightly emphasise that the main beneficiaries of public engagement activities are the members of the public and non-researchers who engage with outreach activities. However, the two-way nature of public engagement is also emphasised in their description: “*engagement is by definition a two-way process, involving interaction and listening, with the goal of generating mutual benefit*”. One way that people can engage deeply with research is by being offered the opportunity to take part in real science experiments. In a commentary in *Journal of Neuroscience*, Heagerty (2015) discusses the “why, when and how” of engaging with the public as (cognitive) neuroscientists and emphasises the importance of sparking dialogue between researchers and non-researchers, rather than seeing the events purely as knowledge dissemination opportunities. With careful planning, it is possible to achieve both remits: by engaging the public in discussion with active research scientists, whilst also capitalising on the opportunity to collect data for scientific projects.

Here we present a case study of a recent public engagement project (“Rhythms of the Brain”), where we aimed to disseminate knowledge about neural oscillations whilst also collecting EEG data to investigate age-related changes in the individual alpha frequency (IAF) and its possible link to fatigue. In the healthy brain, groups of neurons fire together rhythmically (“oscillations”), and these oscillations can be detected using EEG electrodes attached to the scalp. Specific types of oscillations, such as the alpha rhythm (in the 8-12 Hz range), are strongly associated with vision and attention (Thut et al., 2012). Both its prominence and relative ease of detection makes the alpha rhythm an ideal candidate within public engagement contexts. Across the general population, the typical alpha frequency range is around 8-12Hz, although the peak alpha frequency (i.e., the frequency with the highest power) tends to vary across individuals. Regardless a large variation across participants, individual alpha frequency (IAF) has been shown in both cross-sectional and longitudinal studies to gradually change throughout the lifespan (Aurlien et al., 2004; Cellier et al., 2021; Chiang et al., 2011; Cragg et al., 2011; Duffy et al., 1984, 1993; Freschl et al., 2022; Grandy et al., 2013; Klimesch, 1999; Knyazeva et al., 2018; Marshall et al., 2002). Peak occipital alpha frequency is typically slower in young children, at around 6 Hz, and peaking at around 10 Hz in older children and adults (Marshall et al., 2002). The total power of this peak alpha oscillation has also been shown across many studies to decrease with advancing age, both during childhood (Tröndle et al., 2022) and into older adulthood (Whitford et al., 2007). This may reflect a change in white matter integrity and/or loss of grey matter volume throughout the lifespan (Grandy et al., 2013). However, a more recent analytic approach, of dissociating the periodic from aperiodic EEG signal, has shown that older adults may simply experience more broadband 1/f ‘noise’ in their visual systems (Voytek et al., 2015), which may have confounded previous analyses of IAF. In an analysis of 2529 people aged 5-22 years old, Tröndle et al. (2022) found that, after correcting for aperiodic signal, alpha power may in fact *increase* rather than decrease during childhood and adolescence, and decrease between 60-79 years old (Cesnaite et al., 2023). Here we address these questions in a large sample of participants.

The second aim of this study was to investigate whether individual alpha frequency, and alpha power, are linked to self-reported measures of fatigue. It is well established that occipital alpha power increases dynamically during experiments that involve prolonged time-on-task, probably reflective of reduced cortical excitability due to the onset of fatigue (Benwell et al., 2019; Craig et al., 2012; Kasten et al., 2016). Identifying an increased alpha power can also be used as a method of detecting (and alerting individuals to) the onset of transient fatigue in high-risk situations, for example when driving (Schier, 2000). At present it remains unclear whether alpha power and individual alpha frequency are associated with more long-term, tonic reports of subjective fatigue. To explore this question, we administered the Multidimensional Fatigue Inventory (MFI; Smets et al., 1995) to participants as they waited to take part in the EEG experiment. The MFI questionnaire is used to quantify the subjective ratings of 5 different fatigue subtypes (general, physical, mental, reduced activity, and reduced motivation) over the preceding few days, and was analysed by inter-correlating each of the subscales against the EEG outcome measures.

In summary, the overarching aim of this study was to replicate well-established findings of age-related changes in occipital alpha frequency and power during the lifespan within a novel, public engagement context. Specifically, we aimed to 1) identify whether individual alpha frequency and power change throughout the lifespan, 2) identify whether individual alpha frequency and power are linked to subjective ratings of fatigue, and 3) assess the overall feasibility of collecting good-quality data for a simple EEG experiment within a public engagement setting. An exploratory analysis of the periodic and aperiodic signal was performed post-hoc, based on the Fitting Oscillations and One-Over-F (FOOOF) algorithm, which was only available after our first wave of data collection had been completed (Donoghue et al., 2020).

## 2. Methods

### 2.1. Participants

A total of 346 participants were recruited (189 female, 156 male, 1 preferred not to say; Figure 1). The mean age was 29.9 years old (range 6-76 years). We aimed to recruit as many participants as possible using convenience sampling, but an *a priori* sample size calculation estimated a minimum sample size of *n* = 191 would allow a small Pearson’s correlation of *r* = 0.2 to be detected between the participant’s age and their individual alpha frequency, with power = 0.8 and alpha = 0.05.

**Figure 1.**
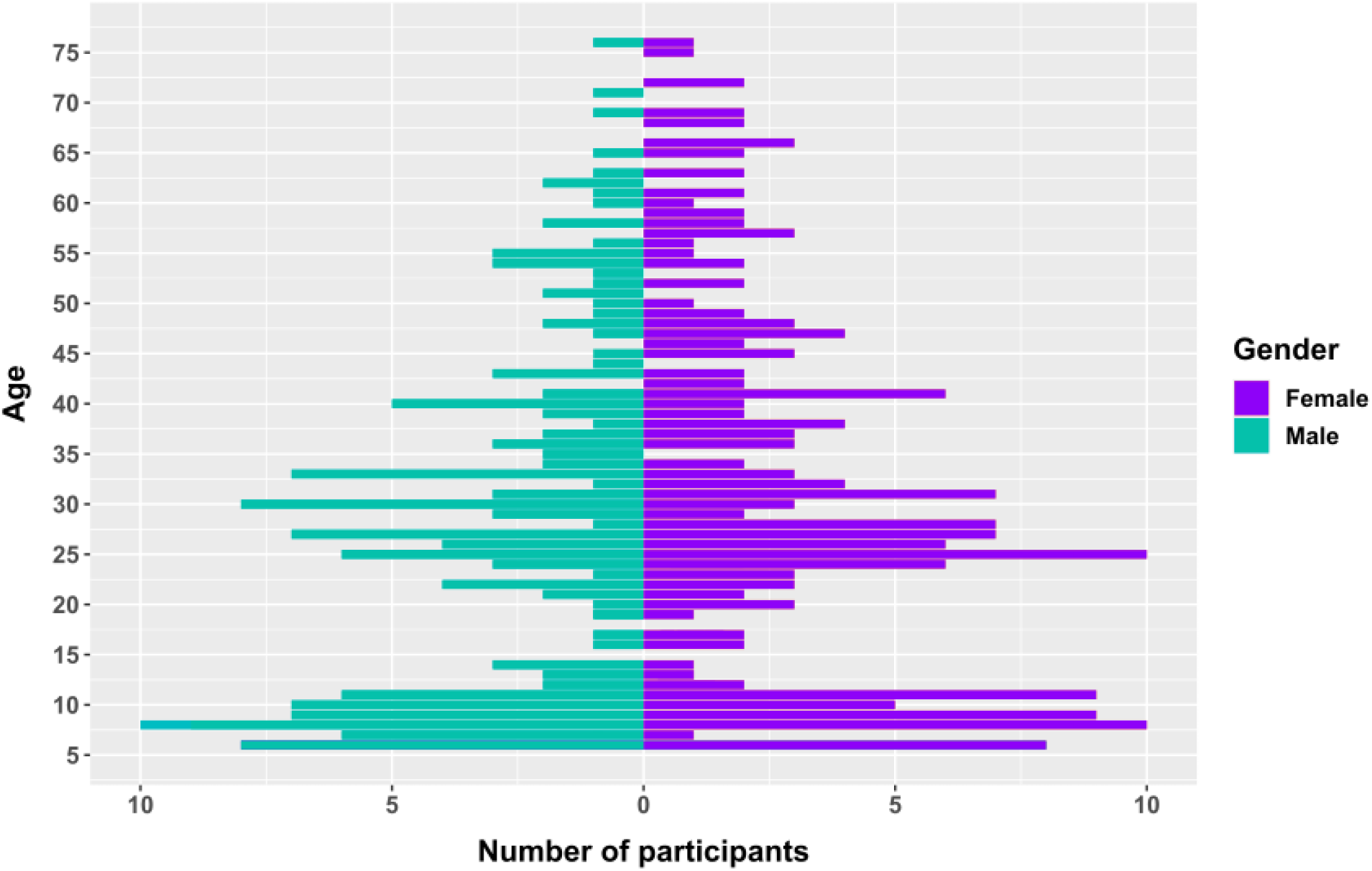
Age and gender distribution of all 346 participants. EEG data were collected from 156 males (indicated by cyan bars) and 189 females (indicated by purple bars) with an average age of 29.9 years.

Data collection took place over 6 days, totalling 29 hours, as part of 2 organised public engagement festivals: Explorathon 2019 and Glasgow Science Festival 2022. Four of the six days were spent at the Riverside Museum, and 2 days at Kelvingrove Art Gallery and Museum in Glasgow, Scotland. Data collection was expected to be completed in 2020 but was interrupted by the pandemic. The only inclusion criterion was a minimum age of 6 years old, with no specific exclusion criteria. The study was approved by the College of Science & Engineering, and the Medicine, Veterinary and Life Sciences ethics committees at the University of Glasgow. All participants formally consented using an electronic tick-box questionnaire, and consent was provided by parents/guardians of children aged under 16 years old.

### 2.2. Procedure

Participants approached our public engagement stall (“Rhythms of the Brain”; see photos in Supplementary Materials) which aimed to engage and educate members of the public on the subject of neural oscillations. They were also invited to “donate their brain waves” as part of a scientific study investigating age-related changes in brain waves and consented to having their signal recorded. If agreed, the EEG electrodes were placed, and they were shown their continuous EEG signal on the laptop screen, then allowed to explore common artifact e.g., eye blinks, and, finally, shown how their alpha rhythms change in size when their eyes are closed compared to when they are open. Each participant was assigned a unique code, and the data were recorded anonymously, with only age and gender recorded. At the end of the session, a debrief form was provided with details of how to withdraw their data if they desired.

### 2.3. Electroencephalography

Two identical BrainVision MR EEG systems were set up, at either end of a table. A single recording electrode was placed on the scalp at the occipital midline. This was identified visually, as being approximately 1 cm above the inion, located between electrode locations Iz (inion) and Oz (occipital midline). SignaGel was used to achieve conductivity between the electrode and the scalp, and the electrode was held in place using an elasticated fabric headband. The ground and reference electrodes were attached to the centre midline of the forehead, approximately 2 cm apart, and held in place using surgical tape. Participants were blindfolded and asked to sit at rest with their eyes closed while the data were recorded for 30 seconds at a 500 Hz sampling rate with an online filter of 0.3 - 100 Hz.

The EEG data were analysed offline using MNE-Python. Since the EEG datasets were of varying lengths, the continuous EEG of all datasets that exceeded 40 seconds were first visually inspected and trimmed to isolate the cleanest 30 second periods (we had aimed to record *around* 30 seconds of eyes-closed data, but some were longer, and these tended to include time periods where the participants were purposefully eliciting eye blinks etc). The datasets that were between 30-40 seconds were not trimmed prior to preprocessing. The resultant signals were bandpass filtered between 4-40 Hz then segmented into 1s epochs. Epochs where the signal exceeded ± 200 µV were removed and the remaining epochs were recombined into a continuous waveform. The spectra of the recombined epochs were calculated using the *welch* function in sciPy (v1.9.3) with a resolution of 0.25 Hz and the following parameters: fs = 500, window = ‘hann’, nperseg = 500, nfft= 2000, detrend = false, return_onesided = true, scaling = ‘spectrum’, average = ‘mean’. The spectra were then decomposed into periodic and aperiodic components using the FOOOF algorithm (Donoghue et al., 2020). The FOOOF algorithm uses a process to fit aperiodic and periodic components to measured power spectra by first flattening the spectra with an initial aperiodic fit, and then identifying peaks in the flattened spectra. The algorithm then uses an iterative approach to refine the aperiodic and periodic fits to create a full model that represents these components separately. The parameters that are used by the algorithm to fit the aperiodic component and identify peaks were set to the following values: peak width limits = [0.5, 12], max number of peaks = infinite, minimum peak height = 0, peak threshold = 2 standard deviations above the mean, and aperiodic mode = ‘fixed’. A detailed explanation of how each of these parameters are used by the algorithm, and more comprehensive overview how the FOOOF algorithm works can be found on the algorithm’s documentation website (https://fooof-tools.github.io/fooof/index.html). Alpha peaks were extracted from the range of 6-15 Hz, due to the anticipated slower peak frequency in young children (Freschl et al., 2022). The total periodic and aperiodic power was obtained by extracting the log(power) value from the *welch*-derived spectrum, at the peak alpha frequency that was obtained using the FOOOF algorithm.

EEG data was recorded from a total of 329 people (*n* = 147 in 2019, and *n* = 182 in 2022). The remaining 17 people who were recruited only completed the MFI questionnaire. Forty participants (12.2%) were excluded post-hoc for one of two reasons: 1) 30 participants (9.1%) had excessively noisy signal, where more than 80% of their segments exceeded 200 µV, and 2) 10 participants (3.04%) had no visible peaks in the 6-15 Hz range. A total of 289 participants (151 female, mean age = 30.1, range = 6-76 years old) were included in the final EEG analysis.

### 2.4. Multidimensional Fatigue Inventory (MFI)

During the first 2 data collection days, participants were also asked to complete the Multidimensional Fatigue Inventory (MFI; Smets et al., 1995), which is a 20-point questionnaire, taking approximately 5 minutes to complete. Each of the 5 subscales is scored between 4-20 points, with higher scores indicating higher levels of fatigue. We were interested in correlating trait fatigue levels, as measured by the MFI, with EEG measures. The MFI was not recorded during the final 4 days of data collection in order to concentrate our resources around collecting EEG. A total of 101 people (56 female, mean age = 34.91, range = 7-69) completed both the MFI questionnaire and EEG recording, and a further 17 people completed only the MFI. Of note, only 3 under 10-year-olds completed the MFI, during which their parents relayed the questions and confirmed that they were able to understand what was being asked.

## 3. Results

All of the raw EEG data and analysis scripts that are used in this article are openly available at https://osf.io/ct2xw/. No withdrawal requests were made following data collection.

### 3.1. Electroencephalography

The mean individual alpha frequency was 9.88 Hz (SD = 1.39, range = 6.08 - 14.97 Hz). There was no linear correlation between age and IAF (*r* = -.018, 95% CI = [-.1, .13], *p* = .77; Figure 2A), but the data were better explained by a loess function which was fit to the data. The peak of the loess curve occurred at 28.1 years old with an IAF of 10.28 Hz. There was a negative linear relationship between the total (unadjusted) alpha power and age, with younger people generally having a higher total alpha power than older people (Pearson’s *r* = -.4, 95% CI = [-.49 -.3], *p* < .0001; Figure 2B). However, there was no correlation between age and *aperiodic-adjusted* alpha power (*r* = -.07, 95% CI = [-.18, .04], *p* = .23; Figure 2C). Both the aperiodic intercept (*r* = -.52, 95% CI = [-.61, -.44], *p* <.0001, Figure 2D) and aperiodic slope (*r* = -.39, 95% CI = [-.48, -.28], p < .0001; Figure 2E) were strongly negatively correlated with age. There were no differences between male and female participants for any of these 5 measures (all *p*-values > .078).

**Figure 2.**
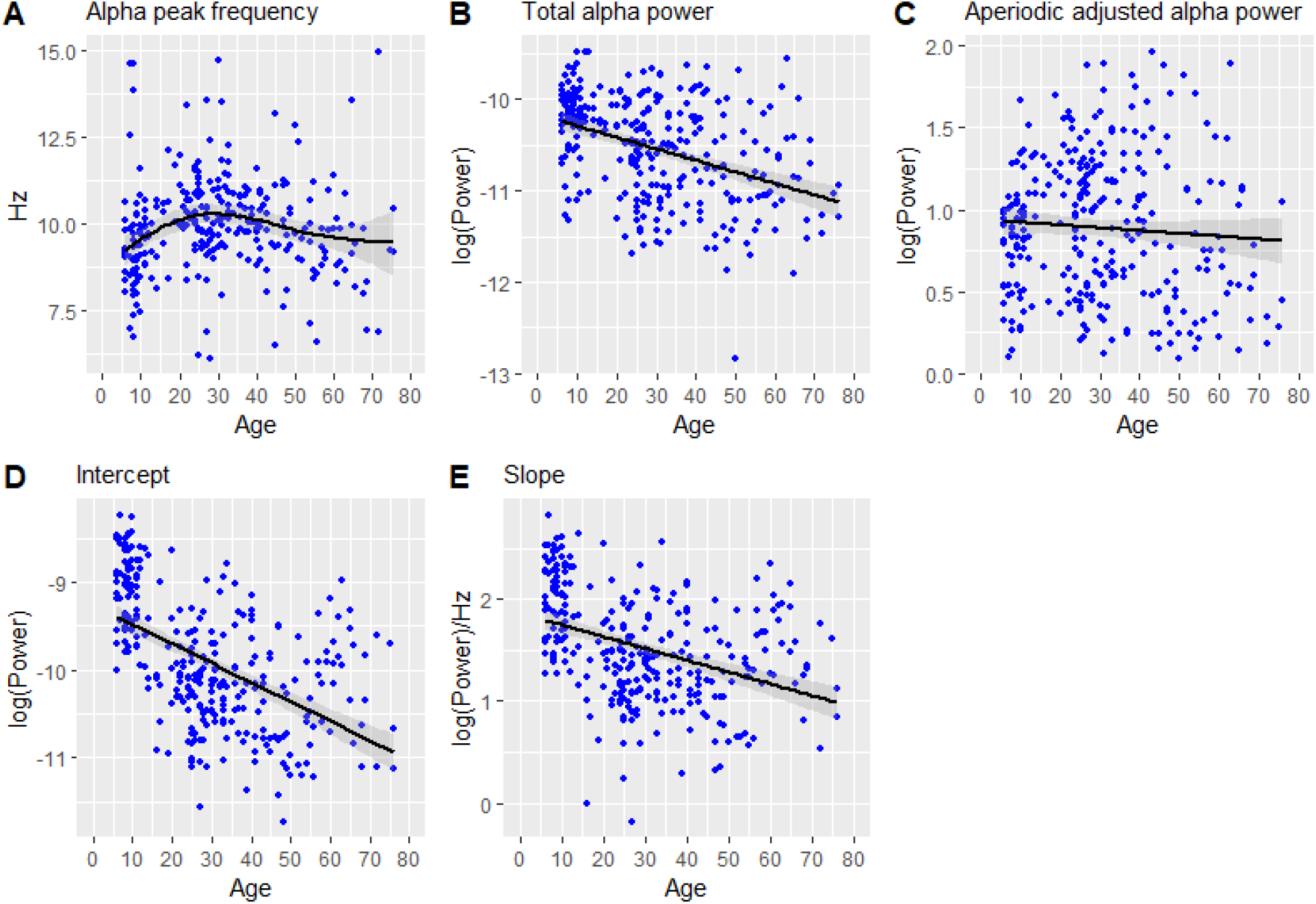
A) Individual alpha peak frequency, B) Total unadjusted alpha power, C) Aperiodic-adjusted alpha power, D) Intercept of the aperiodic slope, E) Slope of the aperiodic exponent. The shaded bands represent the standard error.

The participants were then sorted by age and divided into three bins, each comprising approximately one third of the total number of participants: Age 6-21 (n = 94), age 22-36 (n = 101) and age 37-76 (n = 94) (Figure 3). Splitting the data into three separate bins allows for further comparisons to be made between the age groups, beyond the correlations. Specifically, this allows for a direct comparison of the periodic and aperiodic parameters in the youngest and oldest participants, and mirrors the analysis performed in Tröndle et al. (2022). Of note, the age range of the youngest group in our dataset (ages 6-21) is almost identical to the dataset in Tröndle et al. (2022) (5.04-21.9 years old).

**Figure 3.**
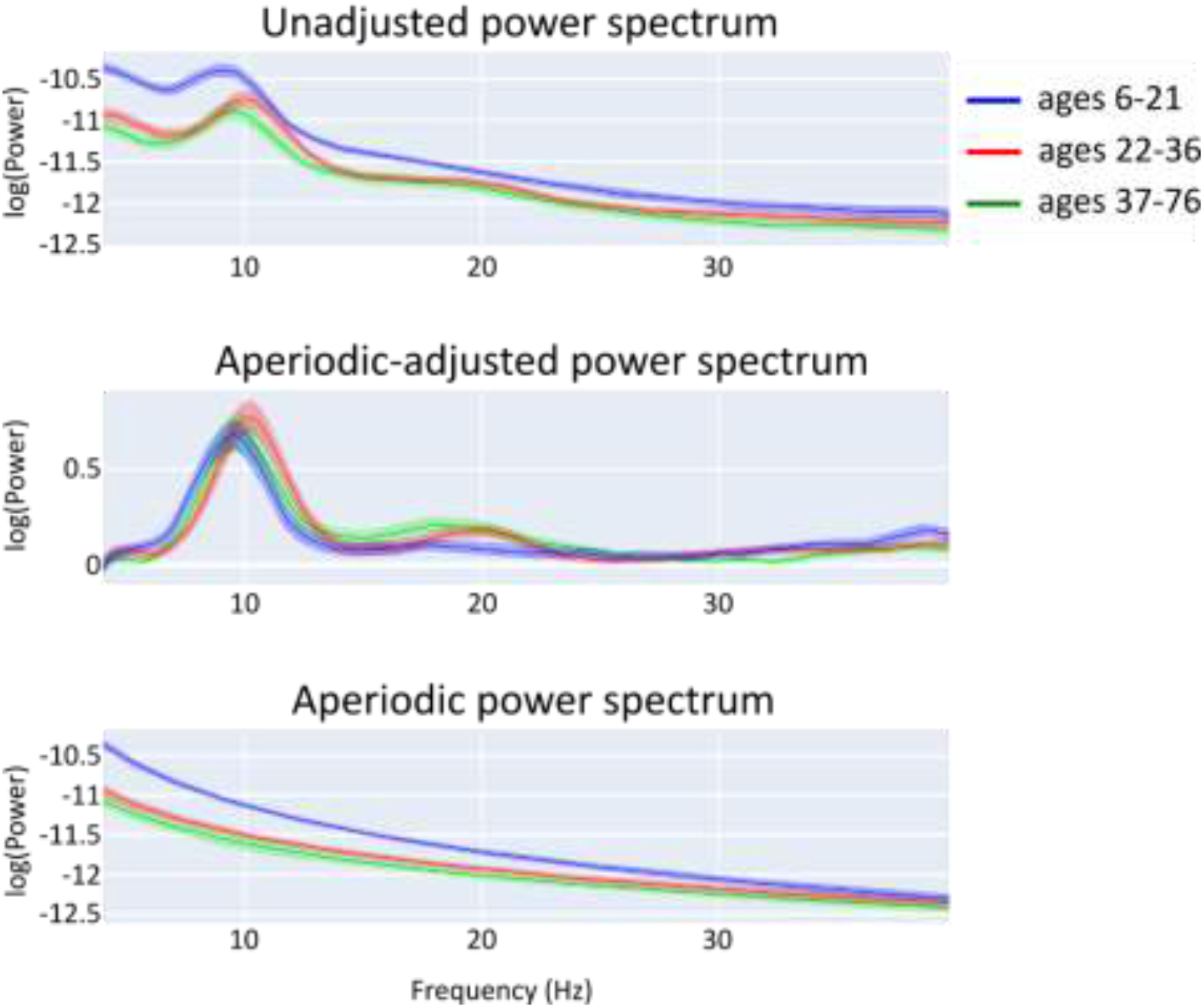
Age-related differences in A) the (aperiodic-adjusted) periodic power spectrum, B) the total measured power spectrum and C) the aperiodic signal. The dataset was divided into 3 bins representing the youngest participants in blue (aged 6-21, n = 94), young adults in orange (aged 22-36, n = 101), and older adults in green (aged 37-76, n = 94). Solid lines represent the mean of each age bin and shaded areas represent the 95% confidence intervals.

Five one-way ANOVAs were then performed, comparing the following 5 EEG outcome measures across the three age bins:

1. *<u>Peak alpha frequency</u>*: There was a main effect of age, *F*(2,286) = 5.85, *p* = .003. Follow-up *t*-tests identified that the middle group (22-36 year olds) had a higher peak frequency than both the youngest group (6-21 year olds; *t*(190) = 3.27, *p* = .0013, *d* = .47) and the oldest group (37-76 year olds; *t*(189) = 2.54, *p* = .012, *d* = .37).
2. *<u>Total (unadjusted) alpha power</u>*: There was a main effect of age, *F*(2,286) = 27.2, *p* < .0001. The youngest group had a higher total alpha power than both the middle group, *t*(190) = 5.69, *p* < .0001, *d* = .81 and the older adults, *t*(163) = 7.1, *p* < .0001, *d* = 1.04. The middle group also had a higher power than the older group, *t*(180) = 2.07, *p* = .04, *d* = .3.
3. *<u>Aperiodic-adjusted alpha power</u>*: There was no effect of age on aperiodic-adjusted alpha power, *F*(2,286) = 1.42, *p* = .24.
4. *<u>Aperiodic slope</u>*: There was a large main effect of age on the aperiodic slope, *F*(2,286) = 48.3, *p* < .0001. The youngest group had a steeper slope than the middle, *t*(181) = 8.51, *p* < .0001, *d* = 1.23 and the older group, *t*(185) = 8.14, *p* < .0001, *d* = 1.19, but there was no difference in slopes between the middle and older groups, *t*(186) = .2, *p* = .84, *d* = .03.
5. *<u>Aperiodic intercept</u>*: There was a large main effect of age on the aperiodic intercept, *F*(2,286) = 96.5, *p* < .0001. The youngest group had a larger intercept than the middle group, *t*(180) = 11.8, *p* < .0001, *d* = 1.7 and the older group, *t*(186) = 11.9, *p* < .0001, *d* = 1.73, but there was no difference in intercepts between the middle and older groups, *t*(178) = 1.48, *p* = .14, *d* = .21.

### 3.2. Multidimensional Fatigue Inventory

The mean score for each of the 5 subscales (where no fatigue = 4 and a high degree of fatigue = 20) was: general fatigue = 11.26, physical fatigue = 9.26, reduced activity = 8.81, reduced motivation = 8.72 and mental fatigue = 10.46. There were no differences between men and women for any subscale (all *t*-values < 1.8, *p* > .074). All 5 subscales were positively correlated with each other, with coefficients ranging between *r* = .67 (between general fatigue and physical fatigue) and *r* = .41 (general fatigue and reduced activity) (Figure 4). Age was positively correlated only with the physical fatigue subtest (*r* = .22, *p* = .016), but was not correlated with mental fatigue (*r* = -.07, *p* = .45), reduced activity (*r* = -.01, *p* = .94), reduced motivation (*r* = .18, *p* = .054) or general fatigue (*r* = .12, *p* = .18). Neither aperiodic-adjusted peak alpha power nor IAF were correlated with any of the 5 subscales (all *r* values < .14 and *r* < .06, respectively). Five separate linear regressions were then performed to assess any interactions between age and IAF, with one of the five MFI sub-scales as the dependent variable in each model, but no interaction was identified (minimum *p* = .35).

**Figure 4.**
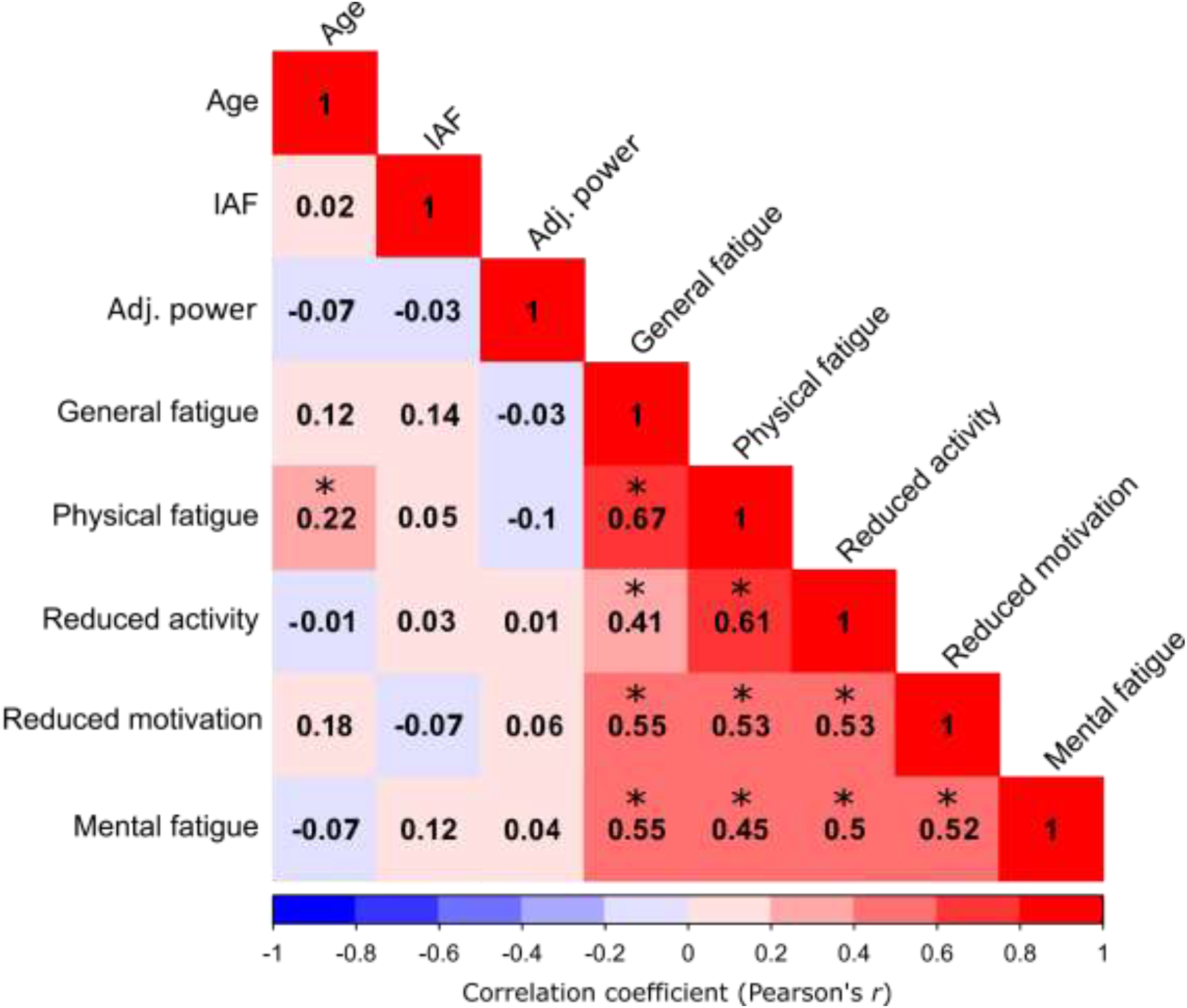
Correlation matrix of the Pearson’s *r* coefficients between age, individual alpha frequency, aperiodic-adjusted alpha power and the 5 MFI subtests. Correlations where *p* < 0.05 are marked with an asterisk. The colour spectrum spans from deep blue, representing a strong negative correlation of - 1, to deep red, representing a strong positive correlation of 1.

## 4. Discussion

We document here our experiences of collecting EEG data, together with questionnaires, within the context of public engagement events. This approach of bringing cognitive neuroscience research equipment out of the lab-based environment and into the community, enabled us to recruit a large sample of participants (*n* = 346), across a wide range of ages (6-76 years old) in a remarkably short period of time (29 hours of testing over 6 days). We confirmed the feasibility of collecting good quality EEG signals outside of the lab, with data from relatively few participants removed from the final analysis due to excessive artifact. Importantly, we successfully replicated previous lab-based findings of a non-linear change in peak individual alpha frequency throughout the lifespan (Aurlien et al., 2004; Chiang et al., 2011; Cragg et al., 2011; Duffy et al., 1984, 1993; Grandy et al., 2013; Klimesch, 1999; Knyazeva et al., 2018; Marshall et al., 2002). We found that individual alpha frequency reached a peak at 28.1 years old (10.28 Hz) and was significantly slower in children and in older adults. The power of the peak individual alpha frequency also appeared to reduce linearly from childhood into older adulthood. However, this correlation was driven by stronger aperiodic signal in children, and there was no observed relationship between age and aperiodic-adjusted alpha power. We did not identify any correlations between the aperiodic-adjusted alpha power and subjective fatigue scores, as measured by the 5 Multidimensional Fatigue Inventory subtests, nor any correlation between the MFI subtests and individual alpha frequency (performed in a subset of *n* = 101 participants).

By decomposing the EEG signal into periodic and aperiodic components (Donoghue et al., 2020), we were able to dissociate the rhythmic brain activity at the alpha frequency from broad-band non-rhythmic activity within the brain. This is an important distinction, because unadjusted alpha power (i.e., including aperiodic signal) is likely to reflect a mixture of different physiological processes and may be misleading when used to link alpha rhythms to specific cognitive states. For example, we were able to replicate previous findings of a decreased aperiodic slope and intercept with increasing age (Tröndle et al., 2022; Cellier et al., 2021; Cesnaite et al., 2023; Hill et al., 2022). The markedly steeper slope in our 6-21 year olds (see Fig 3C) relative to both the 22-36 and 37-76 year olds may reflect a higher prevalence of low-, relative to high-frequency activity in the youngest participants, increased neural “noise” in older age (McIntosh et al., 2010), and/or developmental changes in skull thickness. Similarly, the decrease in the aperiodic intercept during the lifespan may also reflect a generalised reduction in neural activity in older people, although it is important to note that our dataset represents a cross-sectional snapshot of the population, rather than tracking longitudinal changes at the participant level. We cannot exclude the possibility that these differences in the slope and intercept may be spurious and related to differences in drifting eye movements between the groups. We were unable to quantify whether eye movements were present our datasets due to the lack of EOG channels and a measurement of eye movements should be an important quality control to include in future studies.

These aperiodic changes in the EEG signal are apparently distinct from the age-related non-linear increase, then decrease, of the peak individual alpha frequency that we observed. The peak alpha frequency could reflect the speed of sampling of the visual environment (or “temporal resolution”): Samaha & Postle (2015) found that individuals with higher occipital alpha frequencies were better able to identify two flashes, presented with short inter-stimulus intervals, as distinct visual stimuli compared to people with slower individual alpha frequencies. Cecere et al. (2015) present similar results in the audio-visual domain. It may therefore be that the group-level peak alpha frequency identified at 28.1 years old reflects an optimal functioning of the visual system (although see Buergers & Noppeney (2022) for evidence against the influence of trait alpha frequency on perceptual sensitivity). Further, in the absence of repeated, longitudinal recordings to track any shifts in alpha frequency at an individual level during the lifespan, this hypothesis remains an open issue.

Our analyses failed to identify a relationship between alpha power and subjective measures of fatigue. However, this may be related to our choice of questionnaire rather than a lack of relationship between alpha and fatigue *per se*. When completing the Multidimensional Fatigue Inventory, participants were asked to rate their fatigue levels over the preceding few days. In contrast, studies that show a gradual increase in alpha power during the course of an experiment, by way of reduced alertness and increased fatigue with prolonged time-on-task, assess changes in arousal on a more granular scale within the order of minutes (Benwell et al., 2019; Craig et al., 2012; Kasten et al., 2016). The MFI may therefore be an insensitive measure with which to quantify the type of fatigue that is typically associated with fluctuations in alpha power, and a measure that is more sensitive to faster fluctuations in alertness may better reflect the physiological relationship between alpha power and fatigue. Secondly, the overarching concept of “fatigue” encompasses a range of different physiological states, from physical and mental sluggishness to a desire to fall asleep. Given its role in alertness and arousal, we anticipated that any relationship with alpha power would be strongest in the mental fatigue subscale of the MFI (although this was found to be *r* = -.07, *p* = .45), but we also aimed to explore any relationships between alpha power and the other subscales (general, physical, reduced motivation and reduced activity). We found no correlations between any of these, with the largest (although small) effect size of *r* = .18, *p* = .054 associated with reduced motivation.

Given that the recording sessions took place in loud and busy museum environments, we anticipated that the data would exhibit substantially more noise and artifact than an equivalent dataset recorded in the lab. However, only a relatively small number of participants were excluded for this reason. To ensure good quality data, we excluded the participants’ full datasets where the number of noisy segments exceeded 20% of their total recording and only 30/329 participants (9.1%) were excluded for this reason. Although this number of exclusions might represent a large proportion, indeed within the range of the number of the participants who are typically tested within a lab-based experiment (Clayson et al., 2019), in this context where a large sample was tested over a short period of time, it was proportionally relatively few. We have also shown that it is possible to collect questionnaire data during public engagement events, alongside electrophysiological data, to investigate relationships between brain-based measures and self-reported outcomes. However, the experimental design must be carefully considered to fully leverage the opportunities for large-scale data collection that community-based EEG recording can offer. With restricted time windows for data collection per participant, and an additional focus on science communication, experiments must be fast, straightforward and simple, possibly using portable or fully mobile EEG systems.

In terms of participant recruitment, we used a convenience sampling process for this study, by inviting everyone who approached our stall to take part. Although we aimed to recruit a representative cross-section of the population aged 6 years and upwards, there were distinct clusters of participants in the 5-12- and 25-35-year-old age ranges, which tended to represent children, accompanied by their parents. Our stall location within local museums may have contributed to the low number of teenagers taking part (compared to, for example, within a shopping centre or park), and hosting the stall during the week, rather than the weekends, might have increased the recruitment of older adults aged 70+. Due to the constraint of having to sit still with a blindfold for 30 seconds during EEG recording, the minimum age was set to 6 years old, as it was anticipated that children younger than this would not be able to meet this requirement. However, a modified experimental design may have enabled us to collect data from even younger children, assuming that approvals for this had been granted by our local ethics committee. This would have provided a better estimation of the developmental trajectory of alpha rhythms in the very youngest children.

For the sake of simplicity, and to facilitate testing of a large number of people very quickly, we also decided not to collect additional demographic or clinical information from the participants, other than their age and gender. There was no indication that peak alpha frequency or power differed between male and female participants in our dataset, although differences between groups in one, or both, of these measures have been described in previous studies (Cragg et al., 2011; Tröndle et al., 2022). Due to the open nature of our recruitment process, we may have included participants, by design, whose alpha oscillations may be classed as “atypical” relative to the healthy population. For example, there are reports of reduced resting alpha power, resulting in cortical hyperarousal, in people with attention-deficit hyperactivity disorder, schizophrenia and obsessive-compulsive disorder (Newson & Thiagarajan, 2019). Assuming that the appropriate ethical approvals are obtained, more detailed self-reported clinical information about the individual could be collected within public engagement settings to quantify these differences and, based on our collection of *n* = 118 multidimensional fatigue inventory questionnaires in 2 days of testing, other surveys could be administered to isolate other characteristics of the participants, such as personality or other mental states.

Finally, given our experience of collecting data within the community, we have several recommendations and considerations for researchers who wish to use this approach (see Box):

#### Box. Recommendations for community-based EEG recordings

1. **Remember the purpose of public engagement**: Good public engagement is as much about the people, place, methods, aims and impact, as it is about disseminating the results. Its main aim is not purely to disseminate research findings, nor only to collect data, but is a two-way dialogue between researchers and non-researchers. Your activity should primarily focus on engaging your audience, preferably with hands-on tasks (e.g., show them eye blink and muscle artifact from their EEG signal), and data can be collected around this as a secondary objective. Well-planned activities can successfully achieve all of these remits. During this study, participants enjoyed seeing their own brain activity, especially when they could control what appeared on the screen. This generated further questions and allowed them to connect with our research at a deeper level.
2. **Ethical approvals and consent**: Formal ethical approvals must be granted by a research ethics committee prior to collecting data from human participants. This extra workload should be factored into the planning stages of your activity. Each ethics board will provide tailored advice regarding the level of consent that is required. This may be minimal, depending on the type of data that will be collected e.g., The British Psychological Society’s Code of Human Research Ethics states that *“For de-identified-at-source, non-sensitive data, consent may usually be considered to have been given by the act of participation or by ticking a box”* (The British Psychological Society, 2021). Specific care around the issue of informed consent must be taken when collecting data from children and individuals with communication and/or learning difficulties. Debrief letters can be distributed after the activity, including the contact details of the researchers, so that the participant’s data can be rescinded if requested. If any photographs or quotes are recorded from individuals taking part in the activity, ensure that written consent is obtained and clearly state for what purpose and where these will be used (e.g., social media, presentations, newsletters etc).
3. **Where and when will you hold your activity:** The time and location of your activity might be identified by you, or allocated e.g., at a stall during a science fair or festival. These factors are vital in guiding the activity that you will deliver: Who is your audience at this location? Might there be a different audience at the weekend compared to weekdays, and in the morning versus the evening? Is the location loud? Do you have sufficient physical space? Do you need access to power sockets, chairs, washing facilities to clean electrodes? Might it be so busy that you need extra staff? You may also be required to carry out a risk assessment of your activity in advance to identify potential hazards and how you will mitigate them. If working in partnership with a festival, speak with the event organisers early in their planning cycle about your target audience. Guidance on the delivery at appropriate events, venues and time slots, should improve the likelihood of engaging with that group. Data collection over an extended period of time would allow for the identification of under-represented groups and targeting of future activities.
4. **What is your research question and activity:** Simplicity is paramount. Some research questions clearly cannot be answered by collecting data outside of the lab, but others can be addressed with a few modifications to the experimental design and setup. Your task should be quick to set up and to complete, aiming for no more than 5-10 mins per person, or potentially longer if the activity is run as a workshop-style event. Apply the minimum number of electrodes, and record for the fewest number of trials and shortest duration needed to inform your research question. At the same time, bear in mind that reducing the number of electrodes may mean that more care is needed when planning scalp electrode locations and the location of the ground and reference. With a single electrode setup, topographical reconstructions are not possible, and eye movement recordings can be a good compromise to reach a cleaner signal offline. It is best to assume that your participants have little to no background knowledge of your research specialty and therefore the instructions for the task must be easy for your audience to understand, with no scientific jargon. Expect your data to have substantially more noise and artifact than an equivalent lab-based setup, so ensure that you have an objective method of quantifying the quality of the data and be prepared to exclude some participants from analysis. However, the increased availability of participants and the resultant larger sample size can counteract this.
5. **What equipment do you need**: New-generation mobile recording devices are portable by design and are well suited for public engagement events. However, standard EEG systems are also often portable and can be used with care. Research-grade hardware is expensive and fragile, so ensure that it is secure during transportation, storage and during data collection. Older systems that have been retired from the lab are ideal for this reason. It is hard to underestimate the impact of bringing real scientific equipment into a public space as part of the main focus of your activity. People enjoy ‘playing’ with the equipment that researchers use, since most people have limited (or no) access to such equipment after leaving school. Therefore, remember to provide a plentiful supply of consumables e.g., electrodes, connectors, conductive paste, tape, blindfolds etc.
6. **Who is on your delivery team:** Aim to recruit more staff members than you think you need. On a busy day, capacity can soon be overwhelmed when whole families or groups want to take part. You may need an additional, fun activity prepared to entertain those who are waiting in the queue, and someone with good rapport with children can go a long way to easing the pressure. Prior to the activity, ensure that all team members understand the key messages, they can answer simple questions about the theme, and/or a team member with more specialist knowledge is available to continue conversations with interested parties. This is also an excellent opportunity for skill development and improving employability for students and early career researchers who may not want to remain in academia.
7. **Diversity and inclusion:** One of the main benefits of bringing cognitive neuroscience out of the lab and into the “real world” is that it is an opportunity to improve the diversity and representativeness of your research. Consequently, it is important to ensure that your activity is accessible to as many people as possible who wish to take part. Consider whether you would be forced to turn away people wearing a head covering, who use a wheelchair, who have vision or hearing impairments, who speak a different language or who are accompanied by small children and take proactive steps to include everyone who wishes to be involved in your activity. Researchers should be prepared to demonstrate the activity on another member of the team in these cases, and should be prepared to provide information in alternative, accessible formats. Demographic data could also be collected to quantify the improved diversity of your sample.
8. **Closing the loop:** The results of the data collection should be fed back to the participants in some meaningful way to let them know how their data has been used. This can be done directly, using a lay summary, if their contact details are retained, indirectly using a social media hashtag or a dedicated event website, or could form an aspect of your future public engagement activity stall. We intend to disseminate the results and experiences of this study in a future science festival engagement stall. Regrettably, people who participate in scientific studies are frequently not updated on the progress or findings of research studies, although they do feel a sense of ownership of their data. “Closing the loop” in this way can improve the quality of data that is collected and can lead to an improved sense of trust between the public and scientific communities (Long et al., 2017; Purvis et al., 2017).
9. **Be aware of the limitations:** We are keen to emphasise that researchers must carefully reflect upon the potential limitations of using this approach, and the consequences of these limitations for the questions that can feasibly be answered. For example, if scalp topographies are important to the research question, many more electrodes will be required compared to the fast, single-channel setup that we document here. The inclusion of eye electrodes will also allow for better control of eye-related artifacts. However, adding more electrodes will increase the preparation time, and will undoubtedly have a knock-on effect on the number of participants that can be feasibly tested within the available time frame. In short, although recording data in engagement contexts can potentially overcome some of the major issues we face in cognitive neuroscience, in terms of low sample size and poor participant diversity, it is important to be realistic about far this method can go in improving the field of EEG as a whole.

In conclusion, collecting EEG data during public engagement and outreach events can represent a deep way of engaging non-scientists by providing an opportunity to become involved in real science experiments, and meeting researchers who are active in their fields. We have shown that it is feasible to collect good quality cross-sectional data, with outcomes that are similar to those found in lab-based studies, and that a large number of people can be tested within a short period of time. We provide recommendations for other researchers who wish to incorporate EEG data collection into outreach events regarding the planning and delivery aspects of their public engagement activity.

## Acknowledgements

This work was supported by the Wellcome Trust [209209/Z/17/Z]. This project also received funding from the European Union’s Horizon 2020 research and innovation programme under the Marie Skłodowska-Curie grant agreement [No. 794649 awarded to MR, and 897941 to MB].

